# Single-molecule tracking of RNA-DNA hybrid removal enzymes important for lagging-strand replication

**DOI:** 10.64898/2025.12.17.694926

**Authors:** Daniel J. Foust, Frances C. Lowder, Jessica Chung, Julianna R. Cresti, Lieke A. van Gijtenbeek, Lyle A. Simmons, Julie S. Biteen

## Abstract

The formation of RNA-DNA hybrid (RDH) primers by primase is an essential step in the recruitment of DNA polymerase during replication initiation and for the synthesis of each Okazaki fragment on the lagging strand. In addition to primers, RDHs form through misincorporation of ribonucleotides by DNA polymerase during elongation and by formation of R-loops during transcription. R-loops are three-stranded structures that form when the nascent mRNA anneals to the template DNA strand, displacing the complementary DNA strand. The persistence of RDHs is deleterious to genome stability in all cells because they increase susceptibility to mutations, impaired replication fork progression, DNA double-stranded breaks, and genomic rearrangements. In many bacteria, it is well established that components of the replicative DNA polymerase form a macromolecular complex that can be imaged using single-molecule or ensemble fluorescence approaches. The spatiotemporal regulation of proteins involved in RDH removal during lagging-strand maturation is less clear. Here, we study three proteins that are involved in the removal of RDHs from the lagging strand during DNA replication in the Gram-positive bacterium *Bacillus subtilis*: DNA polymerase I (Pol I), FenA, and RNase HIII. We characterized the behavior of each PAmCherry-tagged lagging-strand enzyme in living cells using single-particle tracking photactivated localization microscopy. In this work, we find that all three proteins are highly mobile, suggesting residence times at their target substrates are below our temporal resolution. We also find evidence that Pol I activity is modulated through interaction with the replisome, whereas FenA and RNase HIII are regulated through access to the nucleoid. Our results provide new insight into how enzymes are recruited to resolve RDHs during lagging-strand replication *in vivo*.

**Significance:** RNA-DNA hybrids (RDHs) are essential, transient intermediates in DNA replication, yet their presence significantly increases the susceptibility of the genome to damage. We characterized the single-molecule behavior of three proteins important for processing RDHs in Okazaki fragments in living bacteria. We find that enzyme activity is modulated by access to the replisome and nucleoid. Specifically, we find that DNA polymerase I is preferentially localized to the replisome, while FenA and RNase HIII dwell times at the replisome are very short and below our detection limit. Our work shows that Pol I, FenA, and RNase HIII turn over rapidly in cells, providing new insight into how lagging-strand replication is coordinated *in vivo*.

## INTRODUCTION

RNA-DNA hybrids (RDHs) are nucleic acid intermediates that form *in vivo* during DNA replication and transcription (1-3). During DNA replication, RDHs are formed by primase to begin or “prime” DNA synthesis and by replicative DNA polymerases when they misincorporate single ribonucleoside monophosphates (rNMPs) (1,4). During transcription, RNA polymerase synthesizes mRNA that base-pairs to the template strand, displacing the complementary DNA strand to form a three-stranded structure called an R-loop (5-7). Therefore, some RDHs are essential to prime DNA replication or form during gene transcription, whereas single rNMPs occur as a consequence of high ribonucleoside triphosphate pools during the replication process (8-10). Although some RDHs are essential, they all pose a significant threat to genome integrity if left unresolved, as the 2′-hydroxyl group of the ribose sugar is susceptible to alkaline cleavage, resulting in breakage of the DNA strand and subsequent formation of a 2′, 3′-cyclic phosphate that is recalcitrant to ligation or extension by a DNA polymerase (11). Furthermore, stretches of rNMPs embedded in DNA or R-loops can block replication fork progression, leading to DNA double-strand breaks or mutagenesis (12,13). To address both the necessity for RDHs and to catalyze their removal across systems, from phages to human cells, conserved enzymes are present to remove the RNA portion of an RDH (14–16).

The Endoribonuclease H (RNase H) family is a highly conserved group of enzymes that spans all of biology (15–17). Ribonuclease HI (RNase HI) recognizes stretches of four or more ribonucleotides base-paired to DNA (18,19). Thus, RNase HI is important for the removal of RNA primers at Okazaki fragments and R-loops (20). Although RNase HI is well-conserved, some bacteria, including *Bacillus subtilis*, encode Ribonuclease HIII (RNase HIII) *in lieu* of RNase HI (21). Like RNase HI, RNase HIII has been shown to help with the removal of RNA from Okazaki fragments, with a major role in the removal of persistent R-loops that form during transcription (12,13,22,23). Ribonuclease HII (RNase HII) recognizes both the 2′-hydroxyl on the ribose sugar and the covalent junction between a ribonucleotide and DNA (24–28). Consequently, RNase HII can make an incision 5′ to a single rNMP, initiating the primary pathway for the removal of single ribonucleotide errors (2,29,30). Because RNase HII can recognize any RNA-DNA covalent junction, it has been implicated in the cleavage of the RNA moiety of Okazaki fragments and the re-priming events that occur when replication forks are blocked and restarted by mRNA transcripts (13,23,31,32).

While RNase H enzymes contribute to Okazaki fragment maturation by shortening the RNA primer, 5′ flap endo/exonucleases (FENs) are essential for lagging strand replication as they ensure that all RNA is removed from each Okazaki fragment (33,34). As a result of their essential function, FENs are highly conserved: eukaryotes encode FEN1, which removes RNA primers from Okazaki fragments, and OEX1, which removes R-loops from mitochondrial DNA, while bacteriophages T7 and T4 encode FEN proteins that contribute to phage replication (35–38). For many bacteria, FEN activity is associated with the N-terminal domain of DNA polymerase I (Pol I), while some bacteria, including *B. subtilis*, encode a standalone protein known as FEN or FenA in addition to the Pol I FEN domain (39–41). Therefore, in many different systems, the combination of RNase H incision and FEN excision activities is critical for replication and genome maintenance, with both classes of enzymes contributing to the removal of RDHs from Okazaki fragments.

Several replicative DNA polymerase complex components can be visualized in living cells using fluorescence microscopy, allowing for investigation of the spatiotemporal dynamics during the replication process (42–51). Given that RNase Hs, FEN, and Pol I are important for lagging-strand DNA replication, understanding where and how these proteins localize has been of interest. Previous work studied the localization of *E. coli* RNase HI-YPet, which showed that RNase HI localized to the replisome under normal growth through interaction between RNase HI and single-stranded DNA-binding protein (SSB) (52). Although the K60E mutant of RNase HI maintained wild-type catalytic activity, the mutation obstructed the C-terminal tail of SSB from binding a pocket on RNase HI; this blocked focus formation and ultimately led RNase HI to localize diffusely in cells (52,53). Further genetic experiments suggested that RNase HI K60E impacts R-loop removal but not RNA removal from Okazaki fragments (52). These experiments led to the conclusion that RNase HI localizes to the replication fork in *E. coli* to clear R-loops that are encountered by the replisome, yielding a barrier-free template for replication (52). Therefore, in *E. coli,* it is clear that RNase HI localizes to the replisome, demonstrating that even its R-loop removal activity is localized to the site of DNA synthesis to aid in the replication process.

Ensemble fluorescence measurements of *B. subtilis* Pol I, FenA (ExoR), RNase HII, and RNase HIII have shown that FenA, Pol I, and RNase HII are localized to the nucleoid while RNase HIII is diffuse throughout cells during normal growth (54). Prior work from the same group using photobleaching to achieve single-molecule detection showed that Pol I and FenA were diffusely distributed in cells under normal growth conditions but formed foci at replisome positions following induction of DNA damage with mitomycin C or UV irradiation (55). More recently, however, they reported that only RNase HIII was recruited to the replisome in response to UV-induced damage and that the positioning of Pol I, FenA, and RNase HII was not affected (54). Additionally, they reported evidence of confined diffusion of Pol I, FenA, RNase HII, and RNase HIII near replisomes under normal growth conditions (54). Although these prior studies provide some information on the single-molecule behavior of proteins involved in lagging-strand synthesis, the level of replisome association remains unclear.

In this work, we use single-particle tracking photoactivated localization microscopy (sptPALM) to probe the spatial distribution dynamics of Pol I-PAmCherry, RNase HIII-PAmCherry, and FenA-PAmCherry in live *B. subtilis* cells. The mobilities of tagged Pol I, RNase HIII, and FenA are consistent with freely diffusing proteins, showing no static or reduced mobility state that correlates with the replisome. Analysis of protein heatmaps suggests that changes in growth or DNA replication impact the ability of these proteins to access the nucleoid. Our quantification of protein positioning shows that Pol I is modestly enriched at replisome positions in cells, whereas FenA and RNase HIII are not. Although some proteins involved in replication or repair have been shown to stably associate with replisomes (42,49,52,56–60), our data support the model that FenA, RNase HIII, and, to a lesser extent, Pol I, have sufficient activity towards their appropriate substrates anywhere on the chromosome without the need for concentrated action at the replisome or another specific location under normal growth conditions. Since all three proteins are important for Okazaki fragment maturation (22,23,34,41), our data demonstrate that the turnover of Pol I, FenA, and RNase HIII at the replisome is likely so rapid that enrichment is rarely observed during normal growth or following the inhibition of DNA synthesis. Our studies point to a recruitment mechanism for *B. subtilis* lagging-strand processing enzymes that is different from the more replisome-centric recruitment mechanism employed in *E. coli*.

## METHODS

### Strain construction

We constructed three strains expressing PAmCherry-Pol I, PAmCherry-FenA, and PAmCherry-RNase HIII, respectively. Additionally, each strain expressed DnaX-mCitrine ectopically under control of a xylose-inducible promoter. See the Supplemental Methods for details.

### Imaging sample preparation

We recovered *B. subtilis* cells from 25% v/v glycerol stocks stored at −80 °C, streaked them onto Luria-Bertani agar plates, and incubated them overnight at 30 °C. Cells were transferred to S7-50 minimal media supplemented with 1% w/v arabinose and 0.13% w/v xylose with an initial optical density at 600 nm of around 0.1. Cells were incubated at 30 °C with 200 rpm shaking for 4 – 5 hours until they reached mid-exponential growth phase corresponding to an optical density range of 0.45 – 0.6. For imaging, 2 µL of cells were transferred to 2% w/v agarose pads in the same S7-50 media and covered with a #1.5H thickness cover glass cleaned by plasma etching for 20 minutes in O_2_. Where HPUra or rifampin were used, equal concentrations were added to the agarose as it cooled and to the cells immediately prior to deposition on the agarose pad so that the concentration of the drug would not vary due to absorption into the pad. 40 µM and 15 µM final concentrations were used for HPUra and rifampin, respectively.

### Fluorescence microscopy

We imaged live *B. subtilis* cells on a custom-built optical instrument configured for epifluorescence microscopy. We used 488-nm (Coherent Sapphire 488-50 CDRH) and 561-nm (Coherent Sapphire 561-50 CW CDRH) continuous-wave lasers to excite mCitrine- and PAmCherry-tagged molecules, respectively. A 405-nm laser (Coherent OBIS) was used to photoactivate 0 – 2 PAmCherry-tagged molecules per cell using 100-ms pulses. Laser beams were directed through the back aperture of an Olympus IX71 microscope and focused at the back focal plane of a 100x, 1.45 numerical aperture objective lens (Olympus X-Apo). A dichroic mirror (Chroma ZT488/561rpc) directed collimated light onto the sample. Emitted fluorescence was collected by the same objective, filtered by transmission through the dichroic mirror and a dual-bandpass filter (Chroma ZET488/561m), and directed onto a qCMOS camera (ORCA-Quest, Hamamatsu C15550-20UP) positioned at the image plane of the IX71 tube lens. We used 2 x 2 pixel binning during image acquisition, resulting in a pixel size of 92 x 92 nm^2^ in the object plane.

We captured single imaging frames with 100-ms exposures under 488-nm excitation to visualize mCitrine-tagged DnaX foci. Movies of PAmCherry-tagged proteins were collected under 561-nm excitation with 5-ms exposure times and the camera operating in photon number-resolving mode. Typically, 2 – 3 6000-frame movies were collected per field of view. User-triggered 405-nm photoactivation pulses were periodically deployed during movie acquisition when no single-molecule spots were visible. Average power densities within the full width at half maximum of the Gaussian beam profiles at the sample plane were 15 W/cm^2^ and 300 W/cm^2^ for 488-nm and 561-nm lasers, respectively. For 405-nm photoactivation, 30 W/cm^2^ power density was used at the beginning of the acquisition and was gradually increased up to 175 W/cm^2^ as the pool of photoactivatable molecules was depleted.

### Single-molecule localization and tracking

To detect, localize, and track single molecules, we used Palmari (https://github.com/hippover/palmari), a plugin for the Python-based image analysis software Napari. Detection of single molecules was achieved by applying a Laplacian-of-Gaussian filter with *σ* = 2 to raw movie frames and using a threshold equal to six times the standard deviation of the filtered image pixel values to identify contiguous pixel regions above the threshold as candidate molecules. Candidate molecules were excluded if their equivalent diameter was below 2.5 pixels or above 10 pixels, or if their eccentricity was above 0.9. For subpixel localization, 10 x 10 pixels^2^ particle images were fit with a 2D Gaussian function using GPUfit (61). Tracking was performed using *DiffusionTracker* with parameters: max_diffusivity = 20, max_blinks = 2, and d_bound_naive = 0.1.

### Segmentation

We segmented cells from phase contrast images using Cellpose (62). We used the pretrained model *bact_phase_cp3* to perform an initial segmentation and then manually refined the segmentation where obvious inaccuracies could be seen. For example, when two cells had clearly completed division, but remained in contact, the model occasionally failed to segment them into two cells.

### State Array analysis

We implemented State Array (SA) analysis using the *saspt* package in Python (63). We used a model that accounted for regular Brownian motion and localization errors (RBME). SA analysis estimates the 2D posterior distribution across a fixed array of discrete diffusion coefficients and localization error values. Our arrays consisted of 100 diffusion coefficients ranging from 10^−2^ to 10^2^ µm^2^/s and 30 localization errors ranging from 0 to 100 nm. 2D posterior distributions were marginalized across the localization errors to produce 1D posterior distributions as a function of apparent diffusion coefficient. We limited our analysis to tracks with at least 4 localizations (3 steps) and no more than 16 localizations.

### Heatmaps

We generated heatmaps using the Spideymaps tool in Python as described previously (64). Briefly, grids were generated for each segmented cell using the cell outline determined from the segmented region. Single-molecule detections and pixel areas for grid elements were summed across cells. Fourfold symmetry was applied so that detections and areas for corresponding grid elements in each of the four cell quadrants were also summed. Total counts for each grid element were divided by the total areas to find apparent count densities. Heatmaps were normalized by dividing by the average density across the entire cell so that the displayed values represent densities relative to the density if particles were uniformly distributed.

To capture uncertainty due to cell-to-cell variation, heatmaps were calculated for 1000 bootstrap iterations using sampling with replacement. For each bootstrap iteration, the weighted Pearson correlation coefficient, ρ̂_w_, was calculated against a bootstrap heatmap found for DnaX-mCitrine foci. Weighting was necessary because the Spideymaps heatmaps generate grid elements with unequal areas.

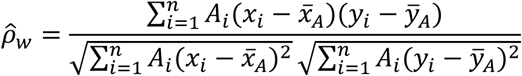

Here, 𝑥_i_ and 𝑦_i_ are the normalized densities for individual grid elements in single-molecule detection heatmaps and DnaX-mCitrine foci heatmaps, respectively. 𝐴_i_ are the areas associated with each grid element. *x̄_A_* and *ȳ_A_* are the area-weighted average densities, by definition, equal to one in the normalized heatmaps.

Heatmaps and Pearson correlation coefficients were calculated for 1000 bootstrap iterations by sampling cells with replacement. Mean heatmap values are displayed. Differences in PCCs were considered statistically significant if the sign of the difference was the same in at least 95% of bootstrap iterations, denoted with *. Similarly, ** and *** denote when the sign of the difference was the same in at least 99% or 99.9% of bootstrap iterations, respectively. “ns” indicates that none of these thresholds were met.

### Radial distribution analysis

Radial distribution analysis is based on the calculation of the ratio between an experimentally measured density, 𝜌_exp_(𝑟), and a null density, 𝜌_null_(𝑟):

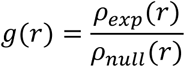

In the context of our experiments, we measured the density of Pol I-PAmCherry, FenA-PAmCherry, and RNase HIII-PAmCherry localizations with respect to DnaX-mCitrine foci. These densities are aggregated across cells. The null density is typically chosen to represent the density if particles are randomly distributed. In bacteria, a common approach to account for how the shape of the cell affects the radial distribution is to simulate randomly placed particles within the cell boundary (65–68). The number of simulated particles is chosen to be equal to the number of experimentally measured particles for that cell so that that cell is appropriately weighted in the aggregate null distribution. We performed a variant of this approach in which we simulated dense rectangular grids of particles uniformly spaced 10 nm apart that filled the bounding boxes surrounding each segmented cell. Grid points falling outside the cell were excluded. We then weighted the contribution of the density calculated for each grid with respect to the DnaX foci by the number of experimental particles measured for that cell. This approach is equivalent to random sampling, but it has the advantage of not requiring repeated sampling to arrive at a stable null distribution.

We calculated the radial distribution functions using radial bins with widths of 100 nm and a maximum radius of 1 µm. To estimate the influence of cell-to-cell variability on the radial distribution functions, 10000 bootstrap iterations were performed by sampling cells with replacement. Differences in radial distribution function amplitudes were considered statistically significant if the sign of the difference was the same in at least 95% of bootstrap iterations, denoted with *. Similarly, ** denotes when the sign of the difference was the same in at least 99% of bootstrap iterations.

## RESULTS

### Single-molecule tracking of lagging-strand, RDH-resolving enzymes in living B. subtilis cells

To directly visualize the subcellular positioning and dynamics of Pol I (*polA*), FenA (*fenA*), and RNase HIII (*rnhC*) with respect to replisomes in live *B. subtilis* cells, we constructed three strains by introducing a C-terminal fusion of the gene for the photoactivatable fluorescent protein PAmCherry (69) to each target protein at its native genomic locus (Figure 1A). Additionally, in each strain, DnaX-mCitrine (70) was expressed ectopically under a xylose-inducible promoter (Figure 1A). DnaX, a subunit of the clamp loader, is commonly used to mark the site of replication (42,71). Accordingly, we imaged DnaX-mCitrine foci to identify the positions of replisomes as described previously (46,72).

**Figure 1.**
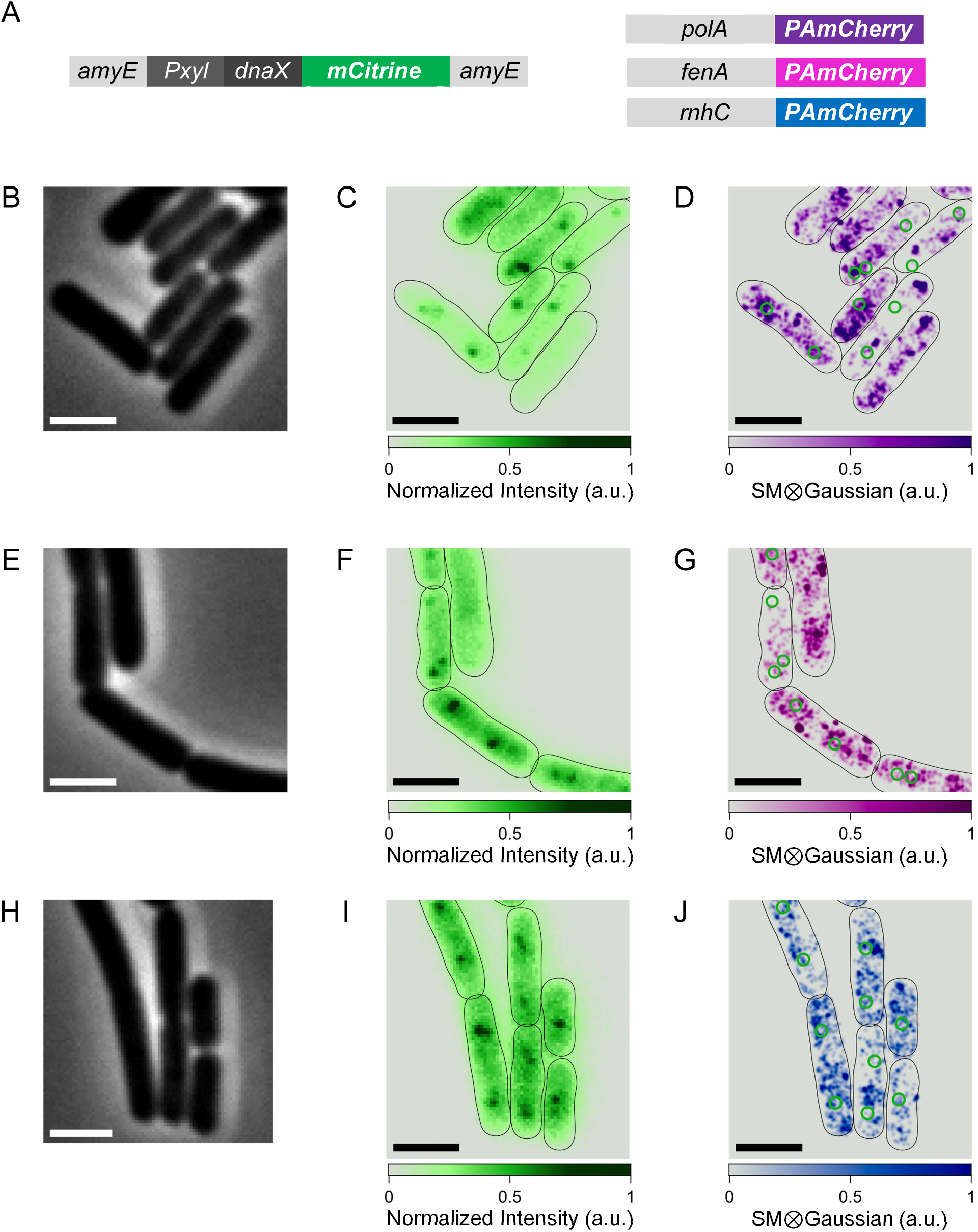
Imaging DnaX-mCitrine and PAmCherry-tagged RDH replication proteins. **(A)** Genetic manipulations introduced to *B. subtilis* strain PY79 to fluorescently label the replisome component DnaX and a ribonuclease with mCitrine and PAmCherry, respectively. **(B)-(D)** Phase-contrast image, mCitrine fluorescence image, and PAmCherry PALM reconstruction for cells expressing DnaX-mCitrine and Pol I-PAmCherry. **(E)-(G)** Same for cells expressing DnaX-mCitrine and FenA-PAmCherry. **(H)-(J)** Same for cells expressing DnaX-mCitrine and RNase HIII-PAmCherry. Black lines indicate cell boundaries determined by segmentation performed on the phase-contrast images. Green circles indicate the positions of replisomes determined by analysis of DnaX-mCitrine images. Scale bars: 2 µm.

We identified rod-shaped cells by phase-contrast microscopy (Figure 1B, E, H), and we imaged DnaX-mCitrine foci under 488-nm excitation (Figure 1C, F, I). To image single lagging-strand replication molecules, we implemented photoactivation localization microscopy (PALM) (73,74) by pulsing 405-nm light to photoactivate a subset of PAmCherry-tagged proteins. For accurate subcellular localization and tracking, the power density of the 405-nm light was tuned to 30 – 175 W/cm^2^, such that 0 – 2 molecules per cell were activated by each 100-ms pulse. Between each pulse, single molecules were imaged under 561-nm excitation with a 5-ms camera exposure time until photobleached. We used a computational analysis pipeline (see Methods) to detect single molecules and find their subpixel localizations. Localizations in adjacent movie frames were algorithmically linked into tracks for analysis of single-molecule dynamics.

We created PALM super-resolution maps from single-molecule localizations for each lagging strand replication molecule and predominantly observed diffuse patterns of subcellular localization for all three proteins (Figure 1D, G, J). We did not detect preferential localization of the lagging-strand proteins near the replisomes by visual inspection of the PALM maps.

### Dynamics of lagging-strand replication proteins with State Array analysis

We quantified the mobility of the PAmCherry-tagged lagging-strand proteins using State Array (SA) analysis, a Bayesian nonparametric framework that estimates the posterior probability distribution of apparent diffusion coefficients from single-molecule tracks (63). Analysis of single-molecule tracks is commonly performed with frequentist approaches that assume a small fixed number of discrete diffusive states (75,76). Recently, Bayesian nonparametric approaches have also gained popularity (77–80). Although these techniques allow a variable number of diffusive states, they are similar to frequentist approaches in assuming a small number of diffusive states, each associated with a discrete diffusion coefficient. The apparent diffusivities of mobile proteins across subcellular environments are influenced by several factors, including varying viscosities and physical confinement imposed by membrane barriers or exclusion from phase-separated condensates. Therefore, we prefer SA analysis because it approximates a continuous distribution of apparent diffusion coefficients by estimating the occupancy of a large number of finite diffusive states, thus providing a more complete description of the mobility of molecules in complex cellular environments (63).

Using SA analysis, we recovered similar diffusion profiles for Pol I-PAmCherry, FenA-PAmCherry, and RNase HIII-PAmCherry (Figure 2). Each diffusion profile features a prominent peak near 2 µm^2^/s, with a broad shoulder extending towards lower diffusion coefficients and a sharp drop-off towards higher diffusion coefficients. Under normal growth conditions, we do not observe any prominent, independent peaks in the diffusion profiles at lower diffusion coefficients. Since substrate binding transiently immobilizes molecules, the data suggest that our imaging approach is not sensitive to transient binding events. Given the overwhelming evidence that these proteins are responsible for resolving RDHs *in vivo* at the replication fork or R-loops that form on the chromosome (13,22,23,34,41), we interpret these data to indicate that binding, cleavage, and release happen on timescales below the 5-ms temporal resolution in these experiments. We consider this timescale to be very fast, since *E. coli* Pol I can be tracked using PALM with experiments at 15-ms timescales (81).

**Figure 2.**
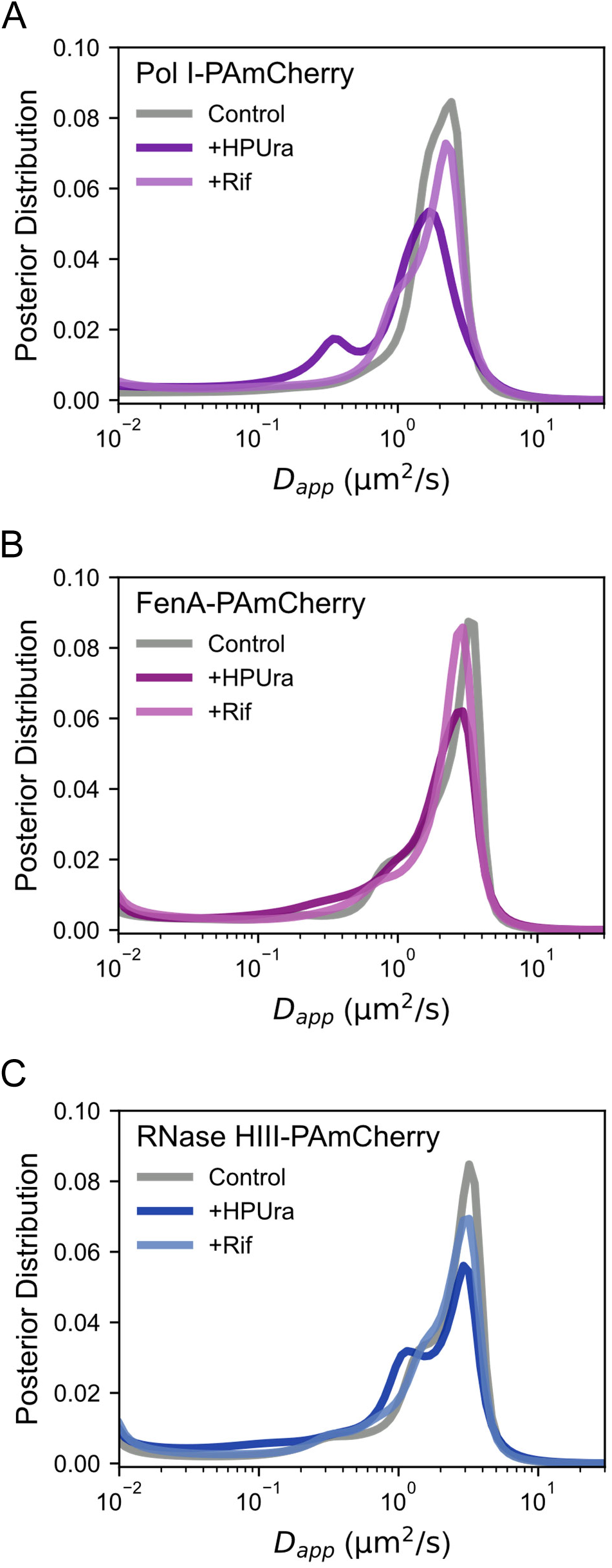
Posterior distributions of apparent diffusion coefficients. Posterior distributions were estimated using State Array (SA) analysis for the following proteins: **(A)** Pol I-PAmCherry, **(B)** FenA-PAmCherry, and **(C)** RNase HIII-PAmCherry.

The sharp drop off to the right of the most probable diffusion coefficient for each protein (Figure 2) is attributable to the limited ability of single-molecule imaging to capture fast-moving molecules (82). Molecules diffusing faster than approximately 2 µm^2^/s are not efficiently detected, even with a 5-ms camera exposure time, due to motion blur. The proximity of the most probable diffusion coefficient for each of our proteins of interest to the detection limit suggests that we sampled subsets of less mobile proteins in our experiments, rather than imaging the complete ensemble of PAmCherry-labeled proteins.

To further our studies, we probed how these lagging-strand processing enzymes respond to inhibition of replication or transcription. First, we used 6-(*p*-hydroxyphenylazo)-uracil (HPUra), which selectively inhibits polymerization by DNA Pol III in Gram-positive bacteria by acting as a dGTP mimic (60,83,84). By inhibiting replication, HPUra reduces the abundance of RNA primers associated with lagging-strand synthesis. Second, we reduced the abundance of R-loop formation with rifampin, which inhibits transcription by blocking the RNAP exit channel for the nascent RNA polymer (85).

Under treatment with 40 µM HPUra, SA analysis indicated shifts towards lower mobility for Pol I, RNase HIII, and FenA (Figure 2). If binding to RNA primers is a significant contributor to the diffusion profiles in control experiments, RNA primer removal would be associated with increased mobility under HPUra-induced depletion. Instead, the reduced RNA primer abundance has the opposite effect (Figure 2). Therefore, the data suggest that interactions with RNA primers do not influence the observed diffusive profiles under normal growth conditions. The modest trend towards low mobility under HPUra treatment may be related to overall changes in cellular viscosity in response to stress (86).

For Pol I and RNase HIII in HPUra-treated cells, we observe the emergence of modest secondary peaks near 0.4 µm^2^/s and 1 µm^2^/s, respectively (Figure 2A, C). To investigate the source of these peaks, we applied SA analysis to the single-molecule tracking data for individual experiments (Figure S1) to determine the variability within the pooled results analyzed in Figure 2. These graphs suggest that the secondary peaks in diffusive profiles for Pol I and RNase HIII under HPUra treatment (Figure 2A, C) are attributable to sample-to-sample variation rather than indicating the emergence of discrete diffusive populations (Figure S1B, H). Notably, 0.4 µm^2^/s and 1 µm^2^/s both correspond to highly mobile molecules; therefore, these peaks do not provide evidence of binding within our temporal resolution. We compared the output of SA analysis to step-size distributions (Figure S2). These distributions agree with the SA analysis as they show modest tendencies towards shorter step sizes in HPUra-treated cells, indicative of reduced mobility (Figure S2). Transcription inhibition with 15 µM rifampin only marginally affected the diffusion profiles of all three proteins (Figure 2). Therefore, interactions with R-loops do not appear to contribute to the measured diffusion of lagging-strand maturation proteins. Altogether, these data suggest that the enzymatic activities of Pol I, RNase HIII, and FenA occur on timescales below the 5-ms camera integration time used in these experiments.

### Subcellular localization of lagging-strand processing enzymes

We observe DnaX-mCitrine foci within both low- and high-density regions of the lagging-strand protein PALM maps, suggesting that apparent colocalization with replisomes is coincidental rather than attributable to dedicated recruitment and prolonged activity at the replication site (Figure 1). To quantify the propensity of each lagging-strand protein to localize to a specific subcellular region, we generated cellular heatmaps by aggregating the single-molecule PALM data (Figure 1) across many cells (Figure 3). We separated the cells into two groups based on cell length. In cells 3.6 – 6 µm long (Figure 3A-G), replisomes display a strong tendency to be found near the one-quarter and three-quarter positions along the cell length (Figure 3A), which indicates that these cells possess two segregated chromosomes. In cells 2–3 µm long (Figure 3H-N), replisomes tend to be found at midcell (Figure 3H), indicative of a single chromosome. We did not include cells 3 – 3.6 µm in length for this analysis because they exhibit greater heterogeneity in replisome positioning (Figure S3). We quantified the tendency of each protein to colocalize with typical replisome positions by calculating the Pearson correlation coefficient (PCC) with heatmaps of DnaX-mCitrine foci.

**Figure 3.**
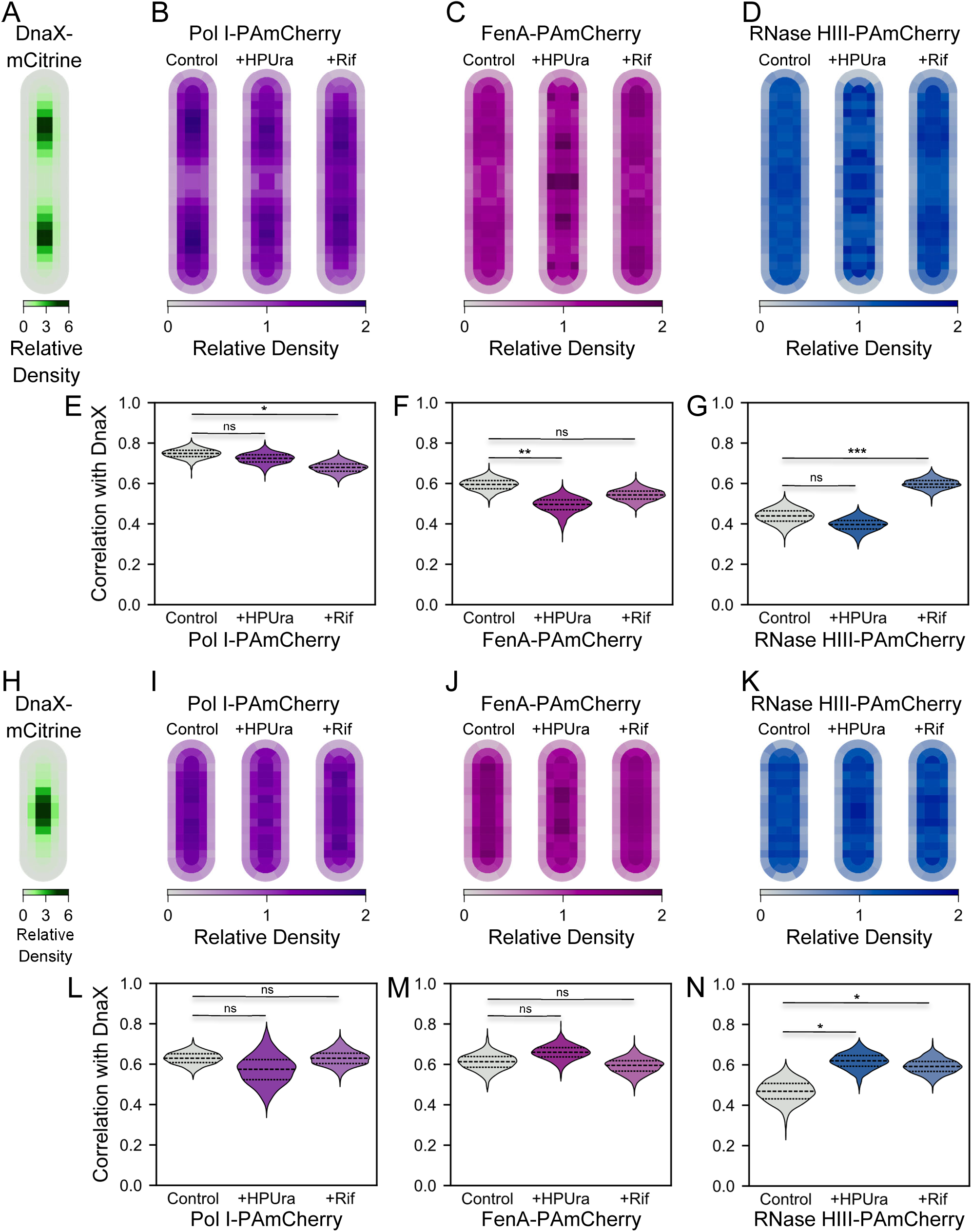
Spatial distributions of PAmCherry-tagged RDH ribonucleases. **(A)-(D)** Heatmaps for DnaX-mCitrine, Pol I-PAmCherry, FenA-PAmCherry, and RNase HIII-PAmCherry, respectively, for cells 3.6 – 6 µm long. **(E)-(G)** Pearson correlation coefficients (PCCs) calculated between the DnaX-mCitrine heatmap in (A) and the RDH ribonuclease heatmaps in (B)-(D). **(H)-(K)** Heatmaps for DnaX-mCitrine, Pol I-PAmCherry, FenA-PAmCherry, and RNase HIII, respectively, for cells 2 – 3 µm long. (**L)-(N)** PCCs calculated between the DnaX-mCitrine heatmap in (H) and the RDH ribonuclease heatmaps in (I)-(K). Heatmaps and Pearson correlation coefficients were calculated for 1000 bootstrap iterations by sampling cells with replacement. Mean heatmap values are displayed. Differences in PCCs were considered statistically significant if the sign of the difference was the same in at least 95% of bootstrap iterations, denoted with *. Similarly, ** and *** denote when the sign of the difference was the same in at least 99% or 99.9% of bootstrap iterations, respectively. “ns” indicates that none of these thresholds were met.

In long cells, Pol I is the most likely protein to colocalize with typical replisome positions (PCC = 0.75; Figure 3B, E). FenA also has a high colocalization (PCC = 0.60; Figure 3C, F). On the other hand, RNase HIII associates much less with typical replisome positions (PCC = 0.45; Figure 3D, G). In short cells, Pol I and FenA heatmaps had PCCs of 0.65 and 0.63 with typical replisome positions (Figure 3I, J, L, M), respectively, and RNase HIII had a substantially lower PCC of 0.47 (Figure 3K, N).

The strong association of Pol I with replisome positions supports its primary function, which is to synthesize DNA and displace RNA primers during lagging-strand synthesis in undamaged cells (34,41). The relatively strong correlation between FenA positioning and the replisomes corroborates the importance of its flap endonuclease activity for RNA primer removal during Okazaki fragment maturation (23,40,41). The weaker association of RNase HIII with the replisomes may indicate a less prominent role in Okazaki fragment maturation and a more significant role in R-loop removal, which is supported by genome-wide R-loop pulldown experiments (13).

We also generated heatmaps and calculated PCCs following replication or transcription inhibition with HPUra or rifampin, respectively (Figure 3). The correlation between Pol I localization density and replisome positioning was significantly decreased in long cells treated with rifampin (Figure 3E). This change was driven by increased accumulation of Pol I near midcell, which contrasts with its exclusion from the midcell under normal growth conditions (Figure 3B). We hypothesize that transcription inhibition by rifampin indirectly inhibits the divisome blocking cell division, allowing more Pol I to localize near the midcell. The alternative explanation, that Pol I localization is sensitive to R-loop depletion, is less likely because we do not observe reduced correlation in rifampin-treated short cells, and Pol I is not known to clear R-loops from cells (Figure 3I, L).

FenA exhibits an increased tendency to localize to the midcell and cell poles in long cells treated with HPUra (Figure 3C, D). This shift significantly reduces the correlation with typical replisome positions (Figure 3F), data consistent with FenA playing a role in RNA primer removal during lagging-strand maturation. The reduced correlation was not recapitulated in short cells (Figure 3J, M), suggesting that FenA activity may be growth-cycle-dependent. Alternatively, the short cells may be actively replicating immediately post-division or dormant and therefore represent a more heterogeneous population than long cells, limiting our analysis.

Similar to FenA, the RNase HIII heatmap under HPUra treatment shows an increased tendency for RNase HIII to localize to the midcell and poles in long cells (Figure 3D); however, this shift was not associated with a change in the correlation with replisome positions (Figure 3G). In short cells, RNase HIII density was more strongly correlated with replisome positions after HPUra treatment (Figure 3K), corresponding to increased protein density at the midcell (Figure 3N). This apparent divergent response of RNase HIII to HPUra treatment in long and short cells may indicate that its activity is linked to the cell growth state.

In cells treated with rifampin, RNase HIII is more strongly colocalized with replisome positions in both long and short cells (Figure 3D, G, K, N). These data suggest that RNase HIII localization patterns are linked to transcription and R-loop formation. We interpret this result to mean that reduced R-loop abundance causes RNase HIII dynamics to shift, with RNase HIII localizing preferentially to the replisome.

Generally, the shifts in localization patterns that we measured in heatmaps (Figure 3) are not easily detected by visual inspection of PALM maps (Figure 1D, G, J). Regions with the highest normalized localization densities across all three lagging-strand proteins are at most two-fold the average density across the cell (Figure 3 B-D, I-K), in contrast to replisomes, which are about six times more likely to be found at their characteristic positions than average (Figure 3A, H). Therefore, the shifts in localization densities we observe are relatively subtle, suggesting that turnover of the molecules at the lagging strand occurs very rapidly.

### Radial distribution function analysis of lagging-strand enzymes with respect to replisomes

Although the heatmaps above show strong correlations between the aggregate densities of Pol I, RNase HIII, and FenA and the DnaX foci, they measure correlations between cell ensembles. Positive correlation between these maps is not necessarily due to direct interactions between the lagging-strand enzymes and replisomes. For instance, we measured the 2D positions of molecules from 3D, rod-shaped bacteria, which inherently leads to more localizations at the cell core (Figure 3B-D, I-K), where replisomes tend to be positioned (Figure 3A, H). Another potential source of spurious correlation comes from association with the nucleoid. Proteins that associate broadly with the nucleoid, but not necessarily directly with replisomes, will still be positively correlated with aggregate replisome positioning across cell ensembles (Figure 3A, H).

To directly probe the spatial relationship between lagging-strand proteins and the replisome, we analyzed the radial distribution of single-molecule localizations in each cell (Figure 4). Radial distribution analysis compares the density of single-molecule localizations as a function of distance from the replisome to the expected density if molecules were uniformly distributed within the cell boundary. The ratio of these two densities, *g*(*r*), is the radial distribution function and measures enrichment (*g*(*r*) > 1) or depletion (*g*(*r*) < 1) relative to the average density.

**Figure 4.**
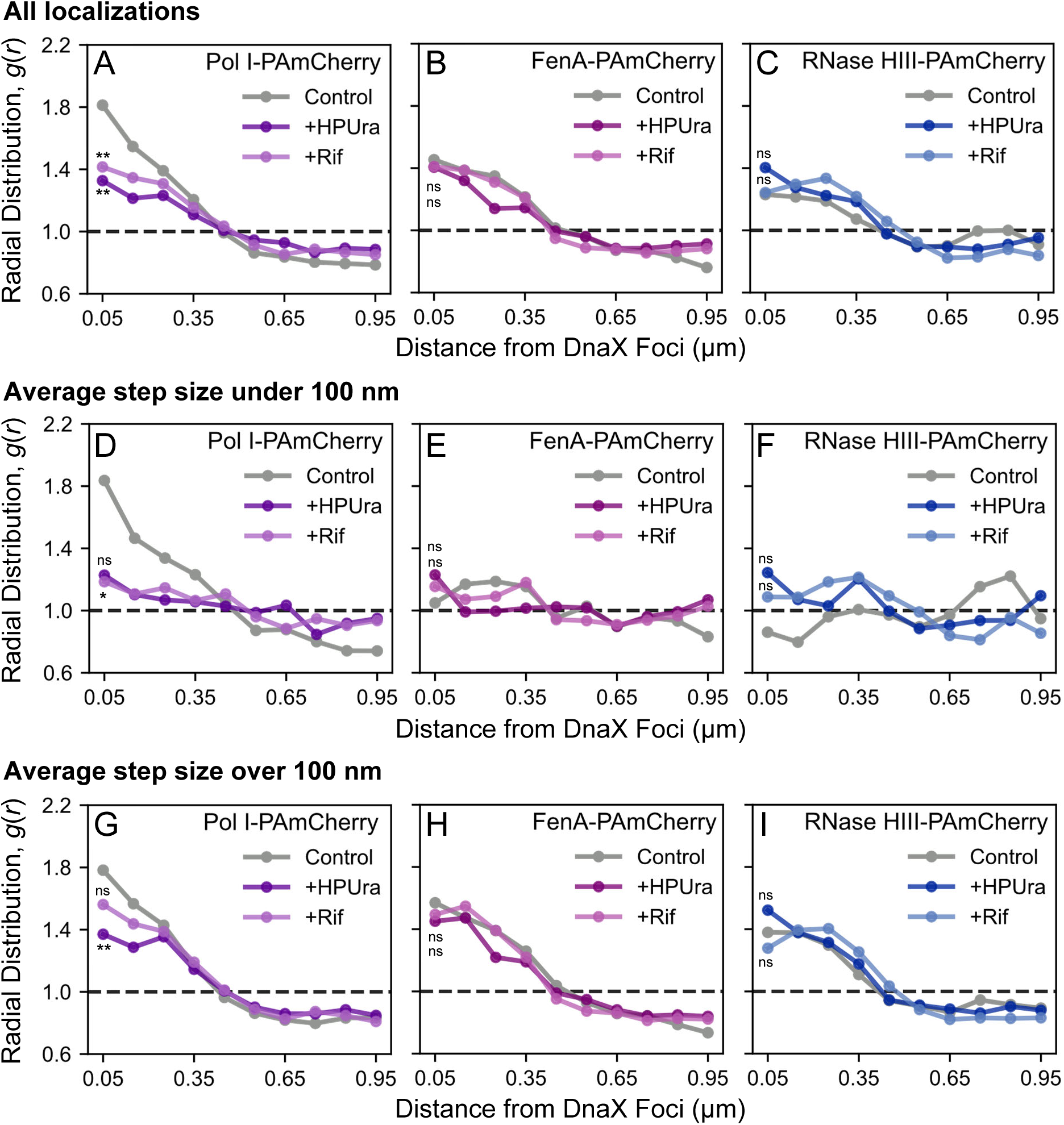
Radial distribution analysis of PAmCherry-tagged RDH ribonucleases. **(A)-(C)** Radial distributions for Pol I-PAmCherry, FenA-PAmCherry, and RNase HIII-PAmCherry localizations, respectively, relative to DnaX-mCitrine foci. **(D)-(F)** Radial distributions for Pol I-PAmCherry, FenA-PAmCherry, and RNase HIII-PAmCherry localizations, respectively, relative to DnaX-mCitrine foci using localizations associated with steps under 100 nm. **(G)-(I)** Radial distributions for Pol I-PAmCherry, FenA-PAmCherry, and RNase HIII-PAmCherry localizations, respectively, relative to DnaX-mCitrine foci using localizations associated with steps over 100 nm. Radial distributions were calculated for 10000 bootstrap iterations by sampling cells with replacement. Median lines are displayed. Differences in radial distribution function amplitudes were considered statistically significant if the sign of the difference was the same in at least 95% of bootstrap iterations, denoted with *. Similarly, ** denotes when the sign of the difference was the same in at least 99% of bootstrap iterations. “ns” indicates that netiher of these thresholds were met.

Pol I, FenA, and RNase HIII are modestly enriched near replisomes, with *g*(*r*) > 1 for smaller *r* (Figure 4A-C). Pol I has the highest and most sharply peaked enrichment near replisomes (Figure 4A). The FenA and RNase HIII radial distributions do not peak sharply near replisomes—rather, they decay gradually and drop below zero near 0.5 µm (Figure 4B, C). Since 0.5 µm is similar to the radius of rod-shaped *B. subtilis* cells, the shape of the radial distributions for FenA and RNase HIII suggests that the apparent enrichment is due to cell shape-related geometric effects and not direct association with the replisome (Figure 4B, C). Conversely, the more sharply peaked radial distribution for Pol I suggests direct association of Pol I with replisomes (Figure 4A). The amplitudes of the radial distributions for Pol I were significantly reduced by HPUra and rifampin treatment (Figure 4A). The radial distributions of Pol I under these conditions (Figure 4A) closely resemble those for FenA and RNase HIII (Figure 4B, C). These data suggest that Pol I association with the replisome is sensitive to inhibition of replication and transcription.

We also considered whether the lagging-strand protein dynamics could be related to replisome proximity. SA analysis of single-molecule tracks for these proteins indicates that the dynamics cannot be attributed to discrete diffusive states (Figure 2). Therefore, we opted to use step sizes to classify detections because single steps provide the most granular temporal view of dynamics (Figure S2). We first classified each detection based on the average size of its adjoining steps (Methods). This classification allowed us to split the localization positions into two groups for separate analysis: those associated with average step sizes below 100 nm, corresponding to lower mobility (Figure 4D-F), and those associated with average step sizes above 100 nm, corresponding to higher mobility (Figure 4G-I). We then applied radial distribution analysis to these subsets of localizations (Figure 4D-I).

For Pol I, the radial distribution for small steps (Figure 4D) has a similar shape and amplitude as the composite of all steps (Figure 4A). The Pol I radial distribution for large steps (Figure 4G) is also similar to the composite Pol I radial distribution (Figure 4A), but its peak is less sharp than in the small steps distribution (Figure 4D). Together, these data support direct interaction of Pol I with the replisome; however, the weak dependence on step size suggests these interactions are transient relative to the 5-ms camera acquisition time.

In contrast to Pol I (Figure 4D), the FenA and RNase HIII radial distributions for small steps are noisy, indistinct, and do not indicate enrichment near replisomes (Figure 4E, F). The corresponding radial distributions for long steps (Figure 4H, I) recapitulate the behavior of the overall radial distributions (Figure 4B, C). Localizations associated with long steps are inherently more strongly constrained by cell geometry because the curvature near cell boundaries confers confinement. Together, these data support that the apparent enrichment of FenA and RNase HIII near replisomes (Figure 4B-C) is attributable to cell geometry effects and not direct interaction with replisomes.

The radial distribution analysis provides important context for the heatmaps and correlation coefficients reported above. This analysis suggests that, for FenA and RNase HIII, the changes in apparent correlations with replisome positioning in response to HPUra and rifampin are likely due to increased or decreased association with the nucleoid rather than direct interactions with replisomes. We conclude that Pol I is associated with replisomes while FenA and RNase HIII are not.

## DISCUSSION

The demonstration that replisomes localize to distinct positions in cells changed the view of how replication occurs and how repair proteins are recruited to replisomes within bacteria (42–51,57). In *E. coli*, it has been shown that RNase HI localizes to replisomes, and single Pol I molecules localize to sites of UV damage with residence times of roughly 2 seconds (52,81). Bulk fluorescence imaging and photobleaching-assisted single-molecule microscopy studies aimed to localize lagging-strand, replication-associated proteins in *B. subtilis* have been less clear, with some studies suggesting replisome association (54,55). We implemented single-particle tracking photoactivation localization microscopy (sptPALM) with 5-ms temporal resolution to examine the behavior of Pol I, FenA, and RNase HIII in *B. subtilis* with high sensitivity.

Here, we find that RNase HIII and FenA are diffusely localized rather than co-localized with the replisome (Figure 4B, C). The degree of association of these proteins with the nucleoid is responsive to changes in replication or transcription status (Figure 3), which slows their subcellular mobilities (Figure 2), but these perturbations do not alter the tendency of these proteins to localize specifically to the replisome (Figure 4). We note that analysis of the protein positioning distribution relative to the average cell replisome positions with Pearson correlation coefficients (PCCs) does yield strong correlations (Figure 3), but we attribute these correlations to the general confinement of FenA and RNase HIII to the nucleoid region and not specifically to the replisome. A more rigorous radial distribution test showed that FenA and RNase HIII do not dwell at the replisome (Figure 4). On the other hand, we did observe that Pol I is recruited to the replisome during normal growth but not when DNA replication or transcription are inhibited (Figure 4A). This correspondence was demonstrated by both the ensemble PCCs and the more stringent radial distribution test. Our findings indicate that the transient enzymatic activity of Pol I during lagging-strand synthesis is manifested as a measurable enrichment at the replisome. Again, for FenA and RNase HIII, we were unable to image these proteins at the replisome at levels above chance events. Given their biochemical activities and considering prior work (20,23,34,41), we expect that FenA and RNase HIII do indeed function during Okazaki fragment maturation: FenA significantly contributes to the release of the RNA flap, and RNase HIII plays a less important role in RNA flap release and a greater contribution to R-loop removal in cells (12,13).

Our results largely agree with two recent imaging studies from the same group (54,55). These works reported that Pol I and FenA are more concentrated in nucleoid-associated regions, whereas RNase HIII is distributed more diffusely (Figure 3) (54). The studies fit the step-size distributions to a two-component model and found two mobile populations for each of Pol I, FenA, and RNase HIII (54). Although the authors described the lower mobility populations they found as “slow” or “static”, the corresponding diffusion coefficients of approximately 0.1 µm^2^/s are more consistent with mobile molecules in our experimental setup. With the 20-ms temporal resolution they used, 0.1 µm^2^/s corresponds to an average step size of approximately 100 nm, much larger than the localization precision in typical single-molecule imaging experiments. The nonparametric Bayesian model we employed indicates a broad diffusive profile rather than two discrete states (Figure 2); however, the two approaches generally agree that these enzymes are highly mobile in cells, consistent with a dominance of free diffusion.

A major discrepancy between the current work and their most recent report is that we do not find evidence that FenA and RNase HIII stably reside at replisomes under normal growth conditions, and we find that Pol I only weakly associates with the replisome (Figure 4) (54). However, the authors posited that long-lived residence of these proteins at replisomes is common based on the percentage of cells with at least one track confined near a replisome for at least 80 ms (54). In our view, that metric is not meaningful without making a comparison to the probability of observing that type of track by chance. Conversely, the radial distribution analysis we implemented in the current work makes an explicit comparison to chance events (Figure 4), and this more robust analysis indicates that interactions between Pol I, FenA, and RNase HIII with the replisome are overwhelmingly transient.

Although *B. subtilis* RNase HIII and *E. coli* RNase HI have overlapping substrate preferences and functional roles in resolving RDHs, including R-loops, these bacteria have unique strategies for regulating their spatial distributions. In *E. coli,* RNase HI binds SSB, and this interaction is critical for its recruitment to replication forks (52,53). Furthermore, *E. coli* RNase HI must remain associated with the replisome to clear R-loops as they are encountered during DNA replication (52). In *B. subtilis*, RNase HIII has not been observed within the SSB interactome (87); if RNase HIII acts at replication forks, it is recruited through a different mechanism. Furthermore, since we do not detect a distinct dwell population for *B. subtilis* RNase HIII, our data suggest that RNase HIII can recognize R-loops or Okazaki fragments without enriching at particular locations. Our data also show that, if RNase HIII is recruited to replication forks to act on RDHs in Okazaki fragments or R-loops encountered by the replisome, the recruitment is stochastic and not directed through a protein-binding partner.

With regard to Pol I, the 5′-to-3′ FEN domain of *E. coli* Pol I is very active because Pol I is largely responsible for both RNA primer removal and DNA synthesis during Okazaki fragment maturation (39). While *E. coli* expresses a discrete flap endonuclease, ExoIX, this FEN lacks key active site residues and therefore has minimal contribution to primer removal due to its minimal activity (88). In contrast, *B. subtilis* expresses both Pol I and a highly active, discrete flap endonuclease, FenA (40,41). In *B. subtilis*, FenA or Pol I can be deleted on their own, while the double mutant is synthetically lethal, demonstrating that either FEN is sufficient for Okazaki fragment maturation (40). Since *E. coli* Pol I supplies both the 5’-to-3’ exonuclease and DNA polymerase activity during Okazaki fragment maturation or DNA repair, its dwell time may be prolonged relative to *B. subtilis* Pol I. *B. subtilis* Pol I primarily supplies the DNA synthesis step, with exonuclease activity coming from FenA, perhaps reducing the dwell time of *B. subtilis* Pol I at the replisome. Altogether, these observations suggest that *B. subtilis* has evolved to rely on stochastic search for RDHs by FenA, RNase HIII, and Pol I, whereas *E. coli* has evolved to favor stable association of RNase HI with the replisome.

In addition to the temporal resolution limits of single-molecule tracking, the interpretation of our results is limited by the imprecision of pharmacological agents as tools for probing replication and transcription *in vivo*. Although the mechanisms of action of HPUra and rifampin against DNA Pol III and RNAP, respectively, are well characterized, knowledge of their indirect effects on overall cell physiology is incomplete (89,90), leaving possible alternative interpretations of our results. For example, we counterintuitively observe stronger nucleoid association of RNase HIII under transcription arrest by rifampin (Figure 3D, K). Rifampin blocks RNA synthesis, which reduces R-loop abundance; consequently, RNase HIII is recruited to the nucleoid despite the depletion of its target substrate. This paradox could be explained by RNase HIII recruitment being mediated by transient interactions with stalled RNAP complexes (91), or by increased access of RNase HIII to the nucleoid as a consequence of the chromosomal response to stress. More precise biochemical tools could provide more insight into how Pol I, FenA, and RNase HIII contribute to resolving replication and transcription-related RDHs. Recently, Cas9 nickases were used to introduce site-specific lesions on the leading or lagging strand to probe the distinct responses at the replication fork to these two types of discontinuities (92). A similar approach could be adapted to more specifically probe RDH resolution during lagging-strand replication. Regarding transcription, RNAP mutants that enhance or suppress R-loops could be used to dissect mechanisms underlying their removal (52).

## Author contributions

DJF, LAS, and JSB designed the research. DJF, FCL, JC, JRC, and LAvG performed the research. FCL, JRC, and LAvG constructed strains. DJF and JC performed imaging experiments and analysis. All authors contributed to the analysis and interpretation of the results. DJF and LAS wrote the first draft of the manuscript with contributions by FCL, JRC, and LAvG to the methods section. All authors revised the manuscript.

## Acknowledgments

This work was supported by the National Institutes of Health (NIH) grant R01 GM144731 to JSB and R35 GM131772 to LAS. JRC was supported in part by NIH training grant: Michigan Predoctoral Training in Genetics T32GM007544. JC’s contributions were supported by the National Science Foundation (NSF) REU 2242779. We thank Dr. Ziyuan Chen, Jewel Ashbrook, and Dr. David Fuller for providing experimental perspective from their unpublished pilot studies, and Kyra Sherman and Sarah El-Mohri for providing technical support.

## Declaration of interests

The authors declare no competing interests.

## Notes

### Competing Interest Statement

The authors have declared no competing interest.

