## Supplemental Information for "Single-molecule tracking of RNA-DNA hybrid removal enzymes important for lagging-strand replication"

**CONTENTS**

- **SI Methods**
- **Figure S1.** Posterior distributions of apparent diffusion coefficients for the individual experiment replicates included in Main Text Figure 2
- **Figure S2.** Step size distributions input to the State Array Analysis in Main Text Figure 2 and SI Figure S1
- **Figure S3.** Positions of DnaX-mCitrine foci versus cell length
- **Table S1.** Primers used in this study
- **Table S2.** Strains and Plasmids used in this study
- **Table S3.** Strains used in this study
- **SI References**

**SI METHODS**

*Plasmid Construction*

pFCL37 - *fenA-PAmCherry*

The editing plasmid to incorporate the FenA-PAmCherry chimera protein was constructed using a Gibson assembly of five PCR-amplified fragments. Piece 1, the *fenA* gene and approximately 1500 bp of upstream sequence, and Piece 3, approximately 1500 bp downstream of *fenA*, were amplified from wild-type genomic DNA with primer pair prFCL83/84 or prFCL87/88, respectively. Piece 2, the *PAmCherry* gene and the 8-residue linker, was amplified from LVG152 using the primer set prFCL85/86. Piece 4, the *Cas9* assembly, and Piece 5, the *AmpR* region, were amplified from pLVG03 using primer pairs oPEB232/prFCL59 and prFCL60/oPEB217, respectively. Amplicons were gel-extracted and combined using NEB HiFi assembly mix (NEB #E2621) at 50 ˚C for 1 hour. The reaction products were transformed into chemically competent MC1061 *E. coli*, and the cells were plated on LB plates supplemented with 100 µg/mL ampicillin after a one-hour recovery. The transformants were grown overnight at 37 ˚C. Colonies were screened using colony PCR with primers oJR349 and oJR350, then successful plasmids were miniprepped and confirmed using Plasmidsaurus whole-plasmid sequencing.

pFCL 40 - *polA-PAmCherry*

The editing plasmid to generate the PolA-PAmCherry chimera was constructed using a similar five-piece strategy. Piece 1, the *polA* gene and approximately 1000 bp of upstream sequence, and Piece 3, approximately 1000 bp downstream of *polA*, were amplified from wild-type genomic DNA with primer pair prFCL81/90 or prFCL97/82, respectively. Piece 2, the *PAmCherry* gene and linker, was amplified from LVG152 using the primer set prFCL101/102. Piece 4, the *Cas9* assembly, and Piece 5, the *AmpR* region, were amplified from pLVG03 using primer pairs oPEB232/prFCL59 and prFCL60/prFCL145, respectively. The fragments were assembled and transformed into MC1061 *E. coli*, as described in the previous section. Colonies were screened using prFCL100 and oJR113, and potential plasmids were miniprepped then confirmed using Plasmidsaurus whole-plasmid sequencing.

pLVG20 - *rnhC-PAmCherry*

Plasmid pLVG01 is a 7264-bp derivative pDR111-sfGFP(Sp) [(1); BGSCID = ECE280)] without the *lacI* gene. pDR111-sfGFP(Sp) was amplified with primers oLVGLS024A and oLVGLS024B. The PCR product was gel-purified, cleaved with BamHI, and self-ligated. The plasmid was propagated in *E. coli* MC1061 (LVG035). Plasmid pLVG02 is derived from pLVG01 by removing the *amyE* recombination domains and the ampicillin resistance gene, while maintaining the *oriT* and spectinomycin resistance gene. pLVG01 was amplified using oLVGLS026A and oLVGLS026B as well as oLVG027A and oLVG027B. The two fragments were gel-purified and combined using Gibson assembly (2). Gibson assemblies consisted of 30 – 80 ng of each PCR product and 1X Gibson assembly mastermix (0.1 M Tris pH 8.0, 5% PEG-8000, 10 mM $MgCl_2$, 10 mM DTT, 0.2 mM dNTPs, 1 mM NAD+, 4 units/mL T5 exonuclease, 25 units/mL Phusion DNA polymerase, 4,000 units/mL Taq DNA ligase) in a total reaction volume of 10 – 12 µL. The reactions were incubated at 50 °C for 90 minutes. Gibson-assembled plasmids were transformed into *E. coli* Top10 or MC1061. PCR fragments were routinely obtained using Phusion polymerase (NEB) or Q5 polymerase (NEB) and gel-purified before Gibson assembly.

The remaining plasmid, pLVG02, is a small preparative plasmid that simplifies assembling multiple large fragments for chromosomal replacements described below. The plasmid was propagated in *E. coli* MC1061 (LVG040). Plasmid pLVG02 was amplified using the primer set oLVGLS026A/oLVGLS023B. A DNA fragment containing the *rnhC* gene including 747 bps upstream was amplified from PY79 genomic DNA with oLVGLS049A and oLVGLS049B, whereas a fragment containing 719 bps downstream of *rnhC* was amplified using oLVGLS049C and oLVGLS049D. A fragment containing a flexible linker translating to AGSGGEAE (encoded by GCGGGTTCTGGAGGTGAAGCTGAA) followed by a codon-optimized allele of PAmCherry was amplified from strain LAM380.1 [(3)] using primers oLVGLS028A and oLVGLS028B. The five fragments were assembled using Gibson assembly, creating pLVG09. This plasmid was propagated in *E. coli* Top10 (LVG123). Plasmid pLVG20 with the backbone of pMiniMad (4) was amplified using oLVGLS053A and oLVGLS053B. A fragment containing *rnhC-up:PAmCherry:rnhC-down* was amplified from pLVG09 using the primer set oLVGLS033A/B. The two fragments were assembled using Gibson, creating pLVG20. This plasmid is propagated in *E. coli* Top10 (LVG149).

*Strain Construction*

All *B. subtilis* strains used are derivatives of PY79. In brief, *B. subtilis* strains were made competent by inoculating LB + 3 mM $Mg_2SO_4$ with one colony of the relevant strain and growing the culture in a rolling rack until $A_{600nm}$ ~ 0.7. This starter culture was used to inoculate MD minimal media (1x PC buffer [10x PC buffer: 107 g/L $K_2HPO_4$, 60 g/L$KH_2PO_4$, 10 g/L trisodium citrate • $(H_2O)_5$], 2% glucose, 50 µg/mL tryptophan, 50 µg/mL phenylalanine, 11 µg/mL ferric ammonium citrate, 2.5 mg/mL sodium aspartate, 3 mM $MgSO_4$). Cells were grown for five to six hours on a rolling rack at 37 ˚C, after which they were transformed with the appropriate gDNA or plasmid DNA. Selective plates used were LB supplemented with either 100 µg/mL spectinomycin (spec), 0.5 µg/mL erythromycin (erm), or 12.5 µg/mL lincomycin (linc) as indicated.

FCL125 - *fenA*::*fenA-PAmCherry*

Competent JRR85 was transformed with pFCL37, plated on LB + Spec, and grown overnight at 30 ˚C. Colonies were re-streaked on LB + Spec and again grown overnight at 30 ˚C. To clear out the temperature-sensitive plasmid, colonies were re-streaked onto LB plates and grown for eight hours at 45 ˚C. Loss of the plasmid and modification of the genome were confirmed by streaking the colonies on LB + Spec and LB + Erm and growing the plates overnight at 30 ˚C. Successful transformants, those that lost the Spec resistance of the plasmid and the Erm resistance of the parent strain, were further confirmed by colony PCR using primer pair oJR349 and oJR350.

FCL137 - *fenA*::*fenA-PAmCherry*, *amyE*::$P_{xyl}$*dnaX-mCitrine*

Competent FCL125 was transformed with gDNA from JWS134, plated on LB + Erm, and grown overnight at 30 ˚C. Colonies were re-streaked onto LB + Erm and LB + 1% starch, then grown overnight at 30 ˚C. Potential transformants were assessed for starch hydrolysis, and those that failed to clear the starch were confirmed with colony PCR using the primer pairs oJR349/350 and oJS437/438.

FCL214 - *polA*::*polA-PAmCherry*
Competent JRR43 was transformed with pFCL40, plated on LB + Spec, and grown overnight at 30 ˚C. Colonies were re-streaked onto LB + Spec and grown overnight at 30 ˚C. Colonies were then re-streaked onto LB plates and grown for eight hours at 45 ˚C to promote loss of the temperature-sensitive plasmid. Successful transformation was assessed by re-streaking onto LB + Spec or LB + Erm and growing overnight at 37 ˚C; cells that grew only on the LB plate were further confirmed by colony PCR using primers prFCL100 and oJR113.

FCL215 - *polA*::*polA-PAmCherry*, *amyE*::$P_{xyl}$*dnaX-mCitrine*
Competent FCL214 was transformed with gDNA from JWS134 and grown overnight on LB + Erm at 30 ˚C. Potential colonies were re-streaked on LB + Erm and LB + 1% starch, then grown overnight at 30 ˚C. Colonies that failed to hydrolyze starch were assessed with colony PCR using primer pairs prFCL100/oJR113 and oJS437/438.

LVG152 - *rnhC::rnhC-PAmcherry*
*B. subtilis* PY79 was transformed with pLVG20, and a colony was selected that showed erythromycin resistance, from which strain LVG150 was generated. Serial propagation under non-selective conditions as described in (5) yielded strain LVG152 containing a chromosomal replacement of *rnhC* by *rnhC-PAmCherry*. The replacement was confirmed by Sanger sequencing.

JRC12 - *rnhC*::*rnhC-PAmCherry*, *amyE*::$P_{xyl}$*dnaX-mCitrine*
Competent LVG152 was transformed with gDNA from JWS134 and grown overnight on LB + Erm + Linc at 37 ˚C. Potential colonies were re-streaked on LB + Erm + Linc and LB + 1% starch and grown on the bench at room temperature. Colonies that failed to hydrolyze starch were assessed with colony PCR using primer pairs prJC37/prJC43 and prJC38/prJC39.

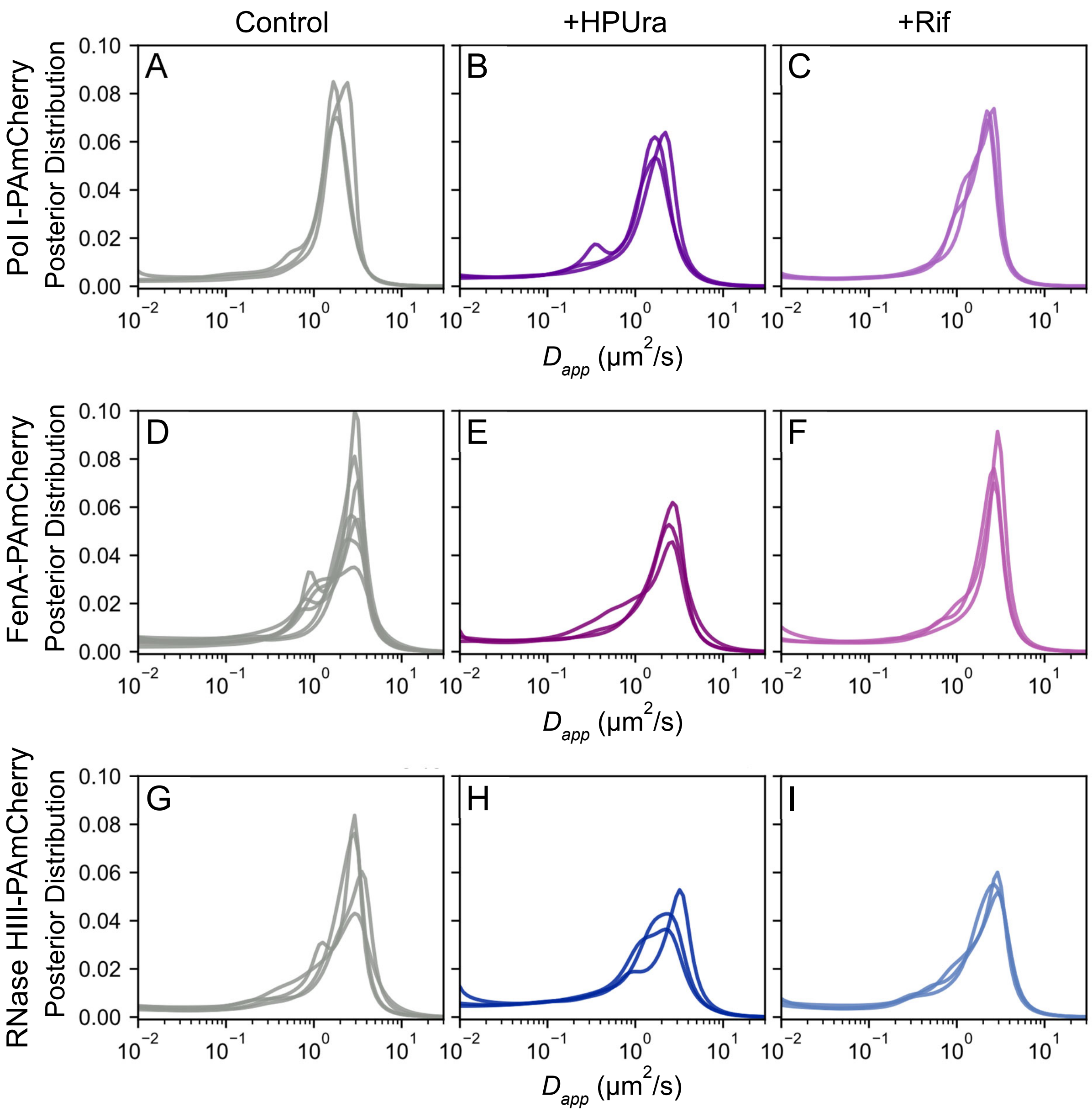


**Figure S1. Posterior distributions of apparent diffusion coefficients for the individual experiment replicates included in Main Text Figure 2.** Posterior distributions were estimated using State Array analysis. **(A)-(C)** Pol I-PAmCherry. **(D)-(F)** FenA-PAmCherry. **(G)-(I)** RNase HIII-PAmCherry.

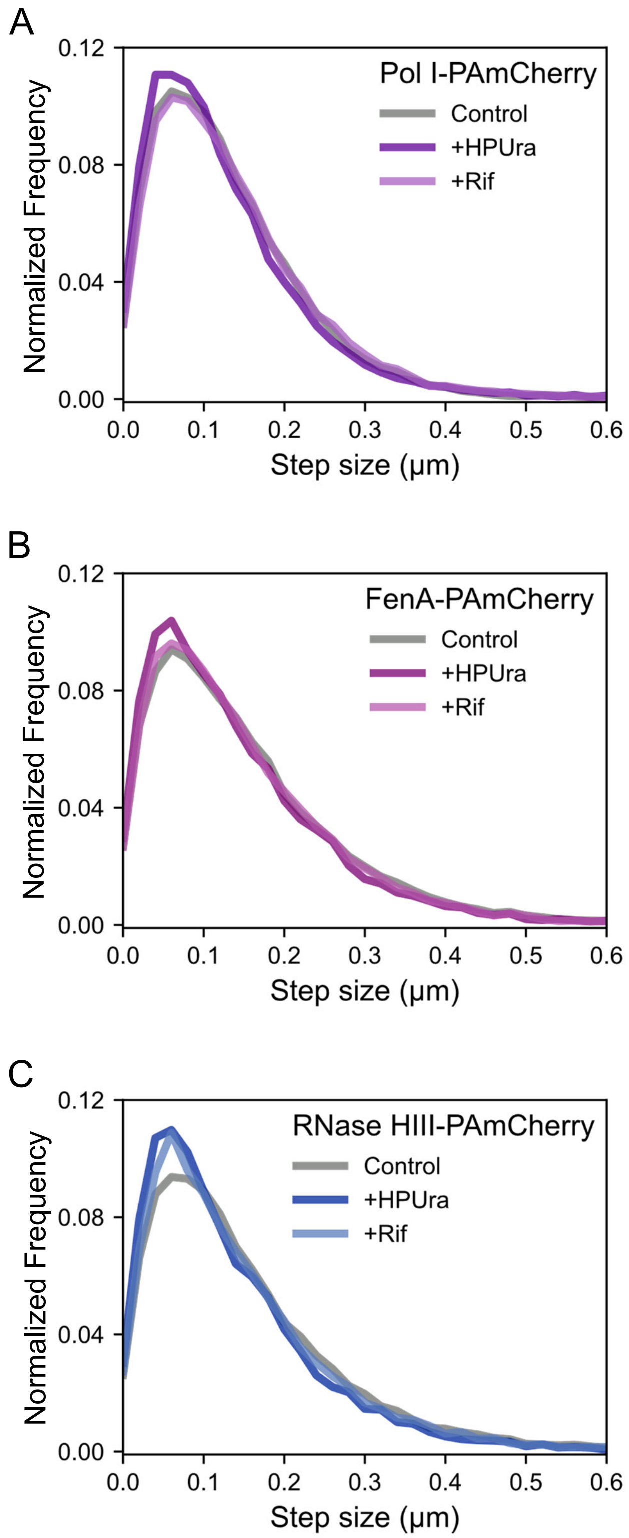


**Figure S2. Step size distributions input to the State Array analysis in Main Text Figure 2 and SI Figure S1. (A)** Pol I-PAmCherry. **(B)** FenA-PAmCherry. **(C)** RNase HIII-PAmCherry.

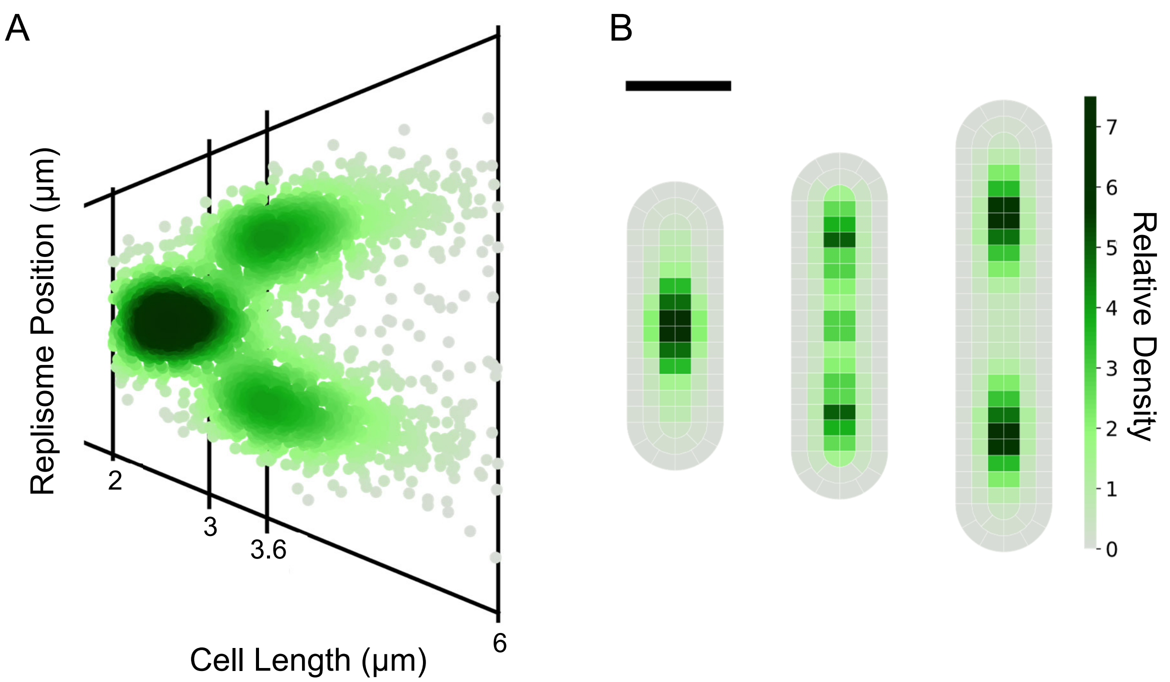


**Figure S3. Positions of DnaX-mCitrine foci versus cell length. (A)** The positions of individual DnaX-mCitrine foci plotted against cell length. **(B)** Heatmaps of DnaX-mCitirine foci corresponding to cell length ranges of 2 – 3 μm, 3 – 3.6 μm, and 3.6 – 6 μm. Scale bar: 1 μm.

**Table S1.** Primers used in this study

| Primer | Sequence | Purpose |
|---|---|---|
| prFCL59 | CTAATCAAGTTTTTTGGGGTCGAGGTGCC GTAAAGCACTAAATCGGAACCCTAAAGG | Amplifying pPB41 *C9* + *SpecR* (R) |
| prFCL60 | CCCTTTAGGGTTCCGATTTAGTGCTTTACG GCACCTCGACCCCAAAAAACTTGATTAGG | Amplifying pPB41 *AmpR* (F) |
| prFCL81 | gcatgctgaattcgtaatgaggttcTATGCTGCGCCG TCTTTATCACGATG | Amplifying *polA* upstream (F) |
| prFCL82 | gcataaccaagcctatgcctacagcTGCTCTTGTATC AGATGTGAAATGC | Amplifying *polA* downstream (R) |
| prFCL83 | catgctgaattcgtaatgaggttcCAGTTCTCGAAAG TAAGAAAACACTTAGACCGG | Amplifying *fenA* upstream (F) |
| prFCL84 | gaaacttcagcttcacctccagaacccgcAACGATCTC TCTAGCGTTCAGC | Amplifying *fenA* upstream (R) |
| prFCL85 | ctgaacgctagagagatcgttGCGGGTTCTGGAGG TGAAGCTGAAGTTTCTAAAG | Amplifying *PAmCherry* + linker (F) |
| prFCL86 | gcaatacaaaacgctagagaggatCTATTTGTAAAG TTCATCCATGCCGCCTG | Amplifying *PAmCherry* + linker (R) |
| prFCL87 | cggcatggatgaactttacaaatagATCCTCTCTAGC GTTTTGTATTGC | Amplifying *fenA* downstream (F) |
| prFCL88 | cataaccaagcctatgcctacagcTCTTCGAACTCT AGCTCCTTCATGTATGG | Amplifying *fenA* downstream (R) |
| prFCL90 | gaaacttcagcttcacctccagaacccgcTTTCGCATC GTACCAAGATGGGCCTGATG | Amplifying *polA* upstream (R) |
| prFCL97 | GGCGGCATGGATGAACTTTACAAATAAAC AGAGATAGGAAGTGATGGATGTGCCGG | Amplifying *polA* downstream (F) |
| prFCL100 | CTTCCGGGTAATGCCGAGG | Colony PCR for *polA* (F) |
| prFCL101 | catcttggtacgatgcgaaaGCGGGTTCTGGAGGT GAAGCTGAAGTTTC | Amplifying *PAmCherry* + linker (F) |
| prFCL102 | ccggcacatccatcacttcctatctctgtttaTTTGTAAAG TTCATCCATGCCGCC | Amplifying *PAmCherry* + linker (R) |
| prFCL145 | gtgataaagacggcgcagcatagaacctcattacgaattca gcatgc | Amplifying pPB41 *AmpR* (R) |
| prJC37 | GTACTGCCTGAACGAGAAGCTATCAC | Colony PCR for *amyE* (R) |
| prJC38 | GTGTCCCATTCAGTGATAAAAG | Colony PCR for *rnhC* (F) |
| prJC39 | CTATTTGTAAAGTTCATCCATGC | Colony PCR for *PAmCherry* (R) |
| prJC43 | GTGAGTTACCAAGCTTTATATCGAGTATTC | Colony PCR for *dnaX* (F) |
| oJR113 | GACAGCAAGCGGGCCG | Colony PCR for *polA* (R) |
| oJR349 | GACGCGAAGCTGAAAAGCTG | Colony PCR for *fenA* (F) |
| oJR350 | GAGGTGAACTAACATGAAAAAGAAAGATA AAG | Colony PCR for *fenA* (R) |
| oJS437 | GGTGGCGGTGGAGGCGGAGGTGGCGGA GGTATGGTGAGCAAGGGCGAGG | Colony PCR for *mCitrine* (F) |
| oJS438 | GGAAGAGTGCGGCCGCCTTATTACTTGTA CAGCTCGTCCATGCCG | Colony PCR for *mCitrine* (R) |
| oPEB217 | GAACCTCATTACGAATTCAGCATGC | Amplifying pPB41 *AmpR* (R) |
| oPEB232 | GCTGTAGGCATAGGCTTGGTTATG | Amplifying pPB41 C9 + SpecR (F) |

| | | |
|---|---|---|
| oLVGLS024A | CCGGGATCCGATGACCTCGTTTCCACCGAATTAGC | Amplifying pDR111 |
| oLVGLS024B | CCGGGATCCGCAGGCCATGTCTGCCCGTATTTC | Amplifying pDR111 |
| oLVGLS026A | CGGTAAGTCCCCAATTAGAATGAATATTTCCC | Amplifying pLVG01 and pLVG02 |
| oLVGLS026B | GATCCATAATGGATTTCCTTACGC | Amplifying pLVG01 |
| oLVGLS027A | GGGAAATATTCATTCTAATTGGGGACTTACCGAAAGAAAC | Amplifying pLVG01 |
| oLVGLS027B | GCCCGTATTTCGCGTAAGGAAATCCATTATGGATCTAGGTGAAGATCC | Amplifying pLVG01 |
| oLVGLS023B | GAGAGTCGAATTCCTGCAGC | Amplifying pLVG02 |
| oLVGLS049A | CCAGGGCTGCAGGAATTCGACTCTCCCAGCATCTGGTTGATTTG | Amplifying *rnhC* |
| oLVGLS049B | CAGCTTCACCTCCAGAACCCGCTGAACGTTTTTTATCAGCAAG | Amplifying *rnhC* |
| oLVGLS049C | GGCGGCATGGATGAACTTTACAAATAGAAAAAAGCTTGCAGATTTC | Amplifying 719 bps downstream of *rnhC* |
| oLVGLS049D | GGATTTCCTTACGCGAAATACGGGCCTGTACGAAGTTGGCTCCG | Amplifying 719 bps downstream of *rnhC* |
| oLVGLS028A | GCGGGTTCTGGAGGTGAAGCTGAAGTTTCTAAAGGCGAAGAAGATAAC | Amplifying PAmCherry |
| oLVGLS028B | TTTGTAAAGTTCATCCATGCCGCC | Amplifying PAmCherry |
| oLVGLS033A | GCATGCTGAATTCGTAATGAGGTTCGGGCTGCAGGAATTCGACTCTC | Amplifying pLVG09 *rnhC*-up:PAmCherry:*rnhC*-down |
| oLVGLS033B | GCATAACCAAGCCTATGCCTACAGCGATTTCCTTACGCGAAATACGGGC | Amplifying pLVG09 *rnhC*-up:PAmCherry:*rnhC*-down |
| oLVGLS053A | GCTGTAGGCATAGGCTTGGTTATGCCGCATCTGTGCGGTATTTCA | Amplifying pMiniMad |
| oLVGLS053B | GAACCTCATTACGAATTCAGCATGCGCCTGGGGTGCCTAATGAGT | Amplifying pMiniMad |

**Table S2.** Strains and Plasmids used in this study

| Plasmid Identifier | Vector | Insert |
|---|---|---|
| pFCL37 | pPB41 + ErmR Protospacer | *fenA-PAmCherry* |
| pFCL40 | pPB41 + ErmR Protospacer | *polA-PAmCherry* |
| pLVG20 | pLVG20 | *rnhC-PAmCherry* |

**Table S3.** Strains used in this study

| Strain Identifier | Relevant Genotype | Citation |
|---|---|---|
| FCL10 | native PY79 | (6) |
| FCL125 | *fenA*::*fenA-PAmCherry* | This work |
| FCL137 | *fenA*::*fenA-PAmCherry*, *amyE*::$P_{xyl}$*dnaX-mCitrine* | This work |
| FCL214 | *polA*::*polA-PAmCherry* | This work |
| FCL215 | *polA*::*polA-PAmCherry*, *amyE*::$P_{xyl}$*dnaX-mCitrine* | This work |
| JRC12 | *rnhC*::*rnhC-PAmCherry, amyE*::$P_{xyl}$*dnaX-mCitrine* | This work |
| JRR43 | *polA*::*ermR* | (7) |
| JRR85 | *fenA*::*ermR* | (7) |
| JWS134 | *amyE*::$P_{xyl}$*dnaX-mCitrine* | (3) |
| LVG040 | *E. coli* MC1061 + pLVG02 | This work |
| LVG123 | *E. coli* top 10+ pLVG09 | This work |
| LVG149 | *E. coli* top 10+ pLVG20 | This work |
| LVG152 | *rnhC*::*rnhC-PAmCherry* | This work |
